# Transcriptome of left ventricle and sinoatrial node in young and old C57 mice

**DOI:** 10.1101/2023.06.16.545319

**Authors:** Jia-Hua Qu, Kirill V. Tarasov, Yelena S. Tarasova, Khalid Chakir, Edward G. Lakatta

## Abstract

Advancing age is the most important risk factor for cardiovascular diseases (CVDs). Two types of cells within the heart pacemaker, sinoatrial node (SAN), and within the left ventricle (LV) control two crucial characteristics of heart function, heart beat rate and contraction strength. As age advances, the heart’s structure becomes remodeled, and SAN and LV cell functions deteriorate, thus increasing the risk for CVDs. However, the different molecular features of age-associated changes in SAN and LV cells have never been compared in omics scale in the context of aging. We applied deep RNA sequencing to four groups of samples, young LV, old LV, young SAN and old SAN, followed by numerous bioinformatic analyses. In addition to profiling the differences in gene expression patterns between the two heart chambers (LV vs. SAN), we also identified the chamber-specific concordant or discordant age-associated changes in: (1) genes linked to energy production related to cardiomyocyte contraction, (2) genes related to post-transcriptional processing, (3) genes involved in KEGG longevity regulating pathway, (4) prolongevity and antilongevity genes recorded and curated in the GenAge database, and (5) CVD marker genes. Our bioinformatic analysis also predicted the regulation activities and mapped the expression of upstream regulators including transcription regulators and post-transcriptional regulator miRNAs. This comprehensive analysis promotes our understanding of regulation of heart functions and will enable discovery of gene-specific therapeutic targets of CVDs in advanced age.

## Introduction

Advancing age is widely recognized as the most important risk factor for cardiovascular diseases (CVDs).^1–4^ Two crucial characteristics of heart function are the rate at which the heart beats and strength of contraction of each beat. Cells within the heart pacemaker, sinoatrial node (SAN), and cells within the left ventricle (LV), are thus crucial to heart functions. SAN pacemaker cells generate electrical impulses that are propagated to other parts of the heart, including LV, in which cardiac myocyte contraction enables LV to pump blood throughout the body.^5–8^ As age advances, across mammalian species from mouse to human, the heart’s structure becomes remodeled, and SAN and LV cell functions deteriorate,^1–4^ increasing the risk for CVDs. Although SAN pacemaker cells and LV myocytes are quite distinct in terms of structure and function,^6, 9^ the different molecular features of age-associated changes in SAN and LV cells have never been compared in omics scale in the context of aging.^1^

In order to gauge the profiles of potential transcriptional mechanisms that changes in SAN and LV in young and old mouse hearts, we applied unbiased discovery transcriptomics to SAN and LV tissues from young adult mice (3-month of age) and those of advanced age (25-26-month of age). Specifically, the transcriptomic analyses employed the deep RNA sequencing (RNA-seq) on four groups of samples, young and old LV (LY and LO, respectively), and young and old SAN (SY and SO, respectively), followed by numerous bioinformatic analyses in order to obtain a comprehensive understanding of transcriptional changes that occur in SAN and LV during aging. In other terms, we sought, not only to define the impact of chamber and age on cardiac gene expression patterns, but also, to define *interactions* between impacts of age- and chamber-specificity on gene expression.

## Materials and Methods

### 1. Animals

Wildtype (WT) C57/BL6 mice were used as the study animals. The 3-month and 26 to 28-month old male mice were included in this project and named as the young and old group respectively. All studies were performed in accordance with the Guide for the Care and Use of Laboratory Animals published by the National Institutes of Health (NIH Publication no. 85-23, revised 1996). The experimental protocols were approved by the Animal Care and Use Committee of the National Institutes of Health (protocol #?).

### 2. Left ventricle (LV) and sinoatrial node (SAN) isolation

LVs and SANs were isolated in the same way described in our previous paper^10^. Briefly, after anesthetization of TGAC8 and WT mice, the hearts were quickly removed. SANs were dissected between inferior and superior vena cava, crista terminalis, and intra-atrial septum and cut into strips perpendicular to the crista terminalis. LVs were dissected from the left chamber of the heart. The isolated SANs and LVs were washed (X3) in the modified low Ca^2+^ Tyrode solution. The isolated LVs and SANs were stored at 4℃.

### 3. RNA extraction, cDNA library preparation, and RNA-seq

By company. In total, we isolated seven tissue samples in each of the four groups (young-LV, old-LV, young-SAN and old-SAN). We sent those samples to DNA Link USA company to conduct the bulk RNA-seq. RNAs were extracted from each sample separately and used as template for cDNA library preparation following the instruction manual of SMARTer Stranded Total RNA-Seq Kit-Pico/Clontech Laboratories, Lnc. The sequencing was conducted on the Illumina NextSeq 500 platform and the data were obtained within the basecalling software RTA2.0.

### 4. RNA-seq data processing

The raw reads were trimmed within Cutadapt and aligned within SAMtools. Using Mm10 as the reference genome, the aligned reads were mapped to calculate the FPKM values in the Cufflinks software set. After quality control and filtration, we remained six young-LV (LY2, LY3, LY4, LY5, LY6 and LY7), six old-LV (LO1, LO3, LO4, LO5, LO6 and LO7), six young-SAN (SY1, SY2, SY3, SY4, SY5 and SY6), and seven old-SAN (SO1, SO2, SO3, SO4, SO5, SO6 and SO7) samples. Keeping genes expressed in all samples, we obtained 9815 genes. The FPKM values were normalized to z-scores for the subsequent hierarchical analysis, principle component analysis (PCA), and ANOVA analysis in the JMP software. Samples were divided into four groups distinctly. The fold-changes and corresponding q-values between two groups were calculated using the Cuffdiff software. The z-score of FPKM values were used to generate heatmaps in RStudio using R language, and the FPKM values were used to generate scatter plots in GraphPad.

### 5. Heatmap

The z-scores of FPKM were imported into RStudio to generate the gene expression heatmap using the pheatmap package in R language. Based on the different expression patterns of the four groups, all available genes were clustered into three clusters. In the uppermost cluster, genes were expressed more in LVs; in the middle cluster, genes were expressed more in SANs; other genes were clustered in the downmost cluster. In addition, the heatmaps for selected family (adenylyl cyclase and surtuin) or class (prolongevity, antilongevity, cardiovascular disease marker, and pre-miRNA) of genes were generated separately.

### 6. GO and KEGG enrichment analyses

Genes in the three clusters mentioned above were used for the GO and KEGG enrichment analyses using the clusterProfiler package in R language. Top ten biological processes (BP), cellular components (CC), molecular functions (MF), and KEGG pathways were displayed in the bubble charts.

### 7. Canonical pathway analysis

All identified genes were uploaded to Ingenuity pathway analysis (IPA) software (QIAGEN, March 2020)^11^. The Ingenuity knowledge base (gene only) about all species and all tissues was selected as the reference set. All Ingenuity supported third party information and Ingenuity expert information were adopted as the data sources, and only experimentally observed relationships were selected to increase confidence. Setting the cutoff as q-value < 0.05 and the absolute value of log2(fold-change) >= 1, IPA recognized: (1) 3333 differentially expressed genes, including 1755 down and 1578 up, within SY vs. LY; (2) 2734 differentially expressed genes, including 1551 down and 1183 up, within SO vs. LO; (3) 127 differentially expressed genes, including 37 down and 90 up, within LO vs. LY; (4) 412 differentially expressed genes, including 263 down and 149 up, within SO vs. SY. The enriched canonical pathways from (1) and (2) were compared to observe the gene expression pattern between the two tissues in young and old mice (Figure 1F).

**Figure 1.**
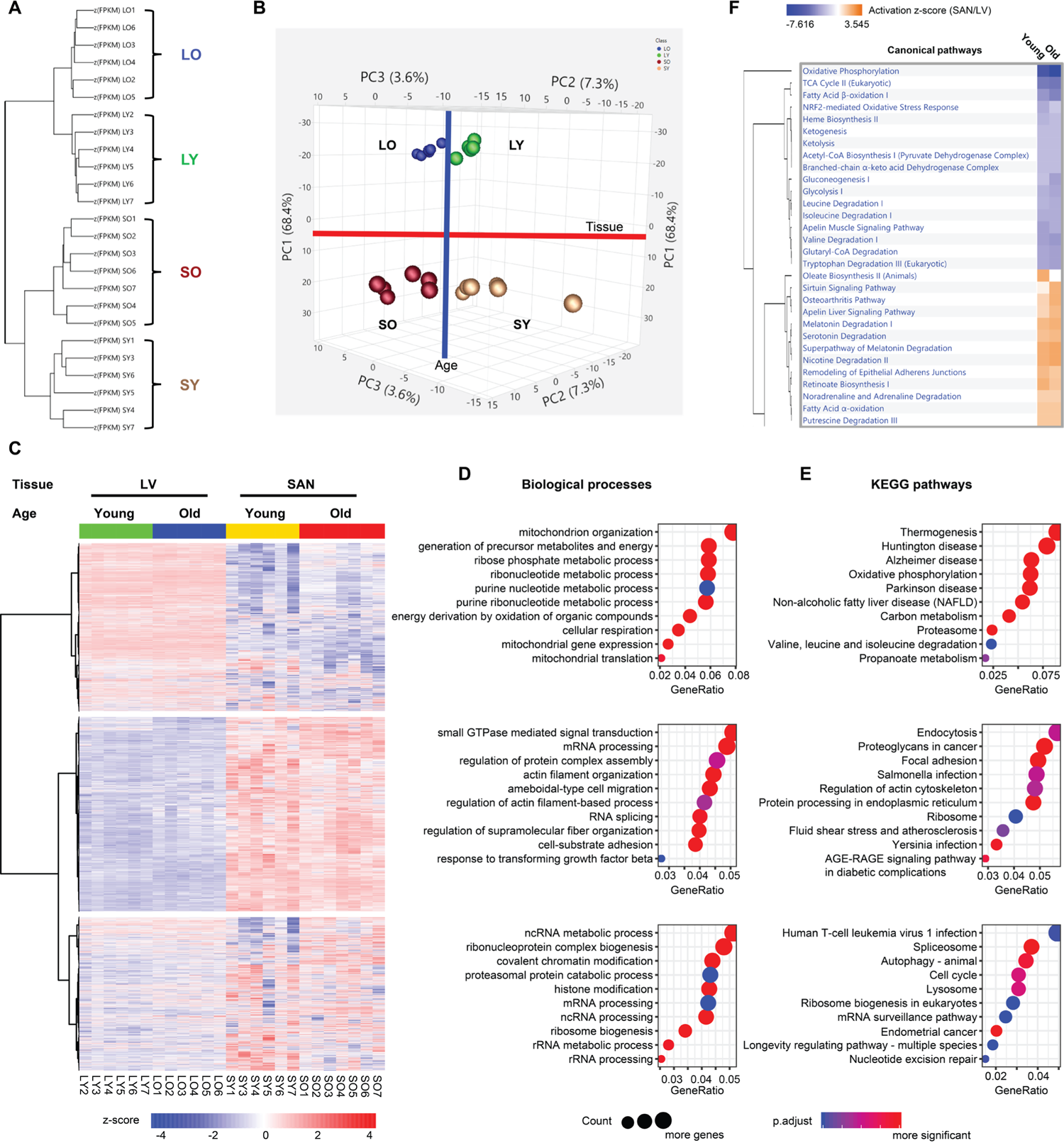
Clustering and enrichment analyses of the four groups of samples. **A.** The hierarchical clustering analysis of the heart tissues. **B.** The PCA of the heart tissues. The LVs (LY and LO) were separated from the SANs (SY and SO) by the first principle component (PC1); the old tissues (LO and SO) were separated from the young tissues (LY and LO) by the second and third principle components (PC2 and PC3). **C.** Heatmap generated with the normalized FPKM values. The genes were clustered into three clusters. **D-E.** Bubble charts of GO biological process analysis (**D**) and KEGG pathway analysis (**E**) with genes in each cluster. The larger the bubble, the more identified genes in the term. The redder, the less adjusted p-value, indicating more significant. **F.** Comparison of the canonical pathways (SAN/LV) enriched within the IPA software. The orange color represents the positive activation z-score and indicates the activated pathway; the blue color represents the negative activation z-score and indicates the inhibited pathway. The deeper the color, the larger the absolute value of the activation z-score. LY: young LV, LO: old LV, SY: young SAN, SO: old SAN.

### 8. KEGG pathway remodeling and visualization

Gene names were transferred into ENTREZIDs preferred by the Pathview online software. The average value of FPKM in each group was calculated and arranged in the order (left to right: LY, LO, SY and SO). After ID transformation and import, there were 9453 IDs recognized by Pathview successfully. The KEGG pathways of cardiac muscle contraction, oxidative phosphorylation, ribosome and protein processing in endoplasmic reticulum, proteasome, and longevity regulating pathway were remodeled and visualized.

### 9. Longevity gene analysis

We mapped the genes identified in our RNA-seq to the GenAge database, a well-admitted longevity gene database, and obtained 85 prolongevity genes and 45 antilongevity genes across the four groups. The comparisons of prolongevity genes and antilongevity genes were separately displayed in heatmaps and volcano plots using R language.

### 10. Upstream transcription regulator analysis

Top 500 genes in the Cluster1 and bottom 500 genes in the Cluster2 were imported into the epigenetic Landscape In Silico deletion Analysis (LISA)^12^ online software to predict the upstream transcription regulators (TRs). In addition, the TRs were also predicted in IPA for the four comparisons in Method 7. Then the results of (1) and (2) were compared, and the results of (3) and (4) were compared in IPA.

### 11. microRNA (miRNA) analysis

In our RNA-seq, the loop pre-miRNAs were detected along with mRNAs. There were eight pre-miRNAs expressed in >= three samples in each group. They were displayed in heatmap to observe their expression across the four groups. There were three significantly differentially expressed miRNAs (q-value < 0.05 and the absolute value of log2(fold-change) >= 1) within LO vs. LY, two within SO vs, SY, ten within SY vs. LY, and 14 within SO vs. LO. The differentially expressed miRNAs were compared and displayed in venn diagrams. In addition, the upstream miRNAs were predicted in IPA for the four comparisons in Method 7. There were two miRNAs predicted to be decreased within LO vs. LY, two miRNAs increased within SO vs. SY, five miRNAs decreased within SY vs. LY, and four miRNAs within SO vs. LO. The regulatory networks for miRNAs to identified genes were generated in IPA.

### 12. RNA extraction, cDNA synthesis, qRT-PCR, and data analysis

The preparation for and performance of qRT-PCR were done as stated in our previous paper in detail^10^. Briefly, RNAs were extracted from isolated mouse LV or SAN with RNeasy Mini Kit (Qiagen, Valencia, CA, United States) and DNAse. 2 mg of total RNA was used for cDNA synthesis with MMLV reverse transcriptase (Promega) in final 50 mL volume. Primers used for each transcript assessed are listed in Supplementary Table (in preparation). qRT-PCR was performed on QuantStudio 6 Flex Real-Time PCR System (Thermo Fisher Scientific) with 384-well platform. Reaction was performed with FastStart Universal SYBR Green Master Kit with Rox (Roche). Normalized to expression of HPRT level, qRT-PCR analysis was performed using ddCt method.

### 13. Statistical analyses

Most statistical analyses were performed using LMP and RStudio (version: 1.1.463) in R language (version: 3.5.3). Some other data processing and statistics were conducted in GraphPad Prism (version: 7) and Microsoft Excel (version: 2019). Adobe Illustrator (version: CC 2019) were also used for graphics without changing data trend or structure. For qRT-PCR analysis, the two-sided unpaired Student’s t-test was used to compare the mRNA expression between two groups, and p-value < 0.05 was taken as statistically significant.

## Results

### 1. Clustering of the age-related transcriptome of LV and SAN

To compare the molecular features between SAN and LV and to reveal the molecular changes in advanced age, we randomly selected 28 heart tissue samples, seven in each group (LY, LO, SY and SO), to conduct the deep RNA-seq. After quality control, six samples in the LY group, six samples in the LO group, six samples in the SY group, and seven samples in the SO group were adopted for subsequent analyses (Supplementary Figure 1). The 25 samples were distinctly separated into four clusters within hierarchical analysis and principal component analysis (PCA), separated by chamber type and age (Figure 1A and 1B). In the hierarchical analysis, the distance between the two chambers was larger than that between different ages (Figure 1A). Consistently, in PCA, SANs were separated clearly from LVs within PC1, accounting for 68.4% difference between samples, and old tissues were separated from young tissues within combined PC2 and PC3 (Figure 1B).

To study the functional characteristics of transcriptional regulations represented in SAN and LV, we displayed the transcriptome in a heatmap and partitioned the genes into three clusters based on their expression patterns across the four sample groups (LY, LO, SY and SO). About a third of the genes were more highly expressed in LV than in SAN and partitioned in Cluster1; another third of the genes were more in SAN than in LV and partitioned in Cluster2; and the remaining third partitioned in Cluster3 (Figure 1C). To confirm the clustering in results, we selected some representative heart marker genes from the literature^5, 7, 13^ and mapped them to our transcriptome. As expected: genes encoding hyperpolarization-activated cyclic nucleotide-gated (HCN) channel isoforms (Hcn1 and Hcn4), T-box transcription factors (Tbx3 and Tbx5), and SPARC-related modular calcium-binding protein 2 (Smoc2) were expressed more in SAN than in LV, and we refer to these as the SAN-specific genes; genes encoding Gap junction proteins (Gja1 and Gjc1), cardiac sodium channel NaV1.5 (Scn5a), and homeobox protein Nkx-2.5 (Nkx2-5) were expressed more in LV than in SAN, and we refer to these as the LV-specific genes (Supplementary Figure 2).

Interestingly, expression of the SAN-specific genes within the SAN was increased with aging, thus increasing difference in expression between the two tissues (SAN vs. LV) in old vs. young mice (the ratio of expression of SAN-specific genes in SO/LO was higher than the ratio in SY/LY). In contrast, with aging, differences in expression of LV-specific genes between the two tissues (LV vs. SAN) became reduced, due to downregulation of these genes in LO or their upregulation in SO (the ratio of expression of LV-specific genes in LO/SO was higher than the ratio in LY/SY) (Supplementary Figure 2B).

### 2. Functional enrichment of each cluster

Next, for each cluster in Figure 1C, we performed GO term (Figure 1D) and KEGG pathway (Figure 1E) enrichment using clusterProfiler package in R language, and canonical pathway enrichment (Figure 1F) using Ingenuity Pathway Analysis (IPA). Pathways enriched within the three clusters of genes differed *mainly* by chamber, regardless of age, but the *degree*, to which some pathways differed by chamber, differed by age.

Functional enrichment analyses showed that **oxidative phosphorylation and metabolism functions** were significantly enriched from those genes in Cluster1 (containing LV-specific genes), as exemplified by: (1) GO biological processes containing mitochondria related processes and various metabolic processes (Figure 1D); (2) GO cellular components of various mitochondrial components (Supplementary Figure 3A); (3) GO molecular functions of electron transfer activity and oxidoreductase activity (Supplementary Figure 3B); (4) KEGG pathways related to thermogenesis, oxidative phosphorylation, carbon metabolism, valine, leucine and isoleucine degradation, and propanoate metabolism (Figure 1E); and (5) canonical pathways related to oxidative phosphorylation, TCA cycle, fatty acid β-oxidation, ketogenesis, ketolysis, acetyl-CoA biosynthesis, and other metabolic pathways (Figure 1F).

In contrast to genes in Cluster1, Cluster2 genes (containing SAN-specific genes) enrichments were more related to **RNA processing, protein processing, and signaling regulation**, exemplified by (1) GO biological processes of small GTPase mediated signaling transduction, mRNA processing, regulation of protein complex assembly, actin filament organization, and RNA splicing (Figure 1D); (2) GO cellular components of extracellular matrix, collagen-containing extracellular matrix, and cytosolic ribosome (Supplementary Figure 3A); (3) GO molecular functions of small GTPase binding, Ras GTPase binding, transcription coregulator activity, GTPase regulator activity, GTPase activator activity, and extracellular matrix structural constituent (Supplementary Figure 3B); (4) KEGG pathways of endocytosis, protein processing in endoplasmic reticulum, and ribosome (Figure 1E); and (5) canonical pathways of sirtuin signaling pathway and noradrenaline and adrenaline degradation (Figure 1F). The longevity regulating pathway was one of the most interesting KEGG pathways that were enriched in Cluster3 (Figure 1E).

## 3. Detailed analyses of functions of clustered genes enriched in GO, KEGG and IPA analyses

### 3.1. Genes linked to energy production related to cardiomyocyte contraction

#### 3.1.1. More highly expressed in LV than SAN

We visualized the KEGG pathway of cardiac muscle contraction to observe the regulation of genes involved in muscle contraction and labeled the average z-score of FPKM of each group (LY, LO, SY and SO) (Figure 2A). As expected, because cardiomyocytes, especially within the LV, contract to maintain the rhythmic blood pressure and provide nutrient and oxygen to the whole body, the expression of genes involved in calcium signaling pathway and contraction (systole and diastole) was higher in LV than SAN. The persistent ion regulation and high-intensive contraction require a large amount of energy. The heart mainly utilizes the energy produced from fatty acid oxidation within the mitochondria. As shown in the illustration of each mitochondrial complex in the KEGG pathway of oxidative phosphorylation, mitochondrial complex genes were more highly expressed in LV than in SAN (Figure 2B). This expression pattern was in accord with the enrichments in mitochondrial components in GO_CC (Supplementary Figure 3A) and mitochondrial processes in GO_BP (Figure 1D), likely linked to a greater energy production required for LV contraction vs. SAN contraction. Compared to LV, SAN displayed higher expression levels of sodium/hydrogen exchanger (NHE) genes (Figure 2A).

**Figure 2.**
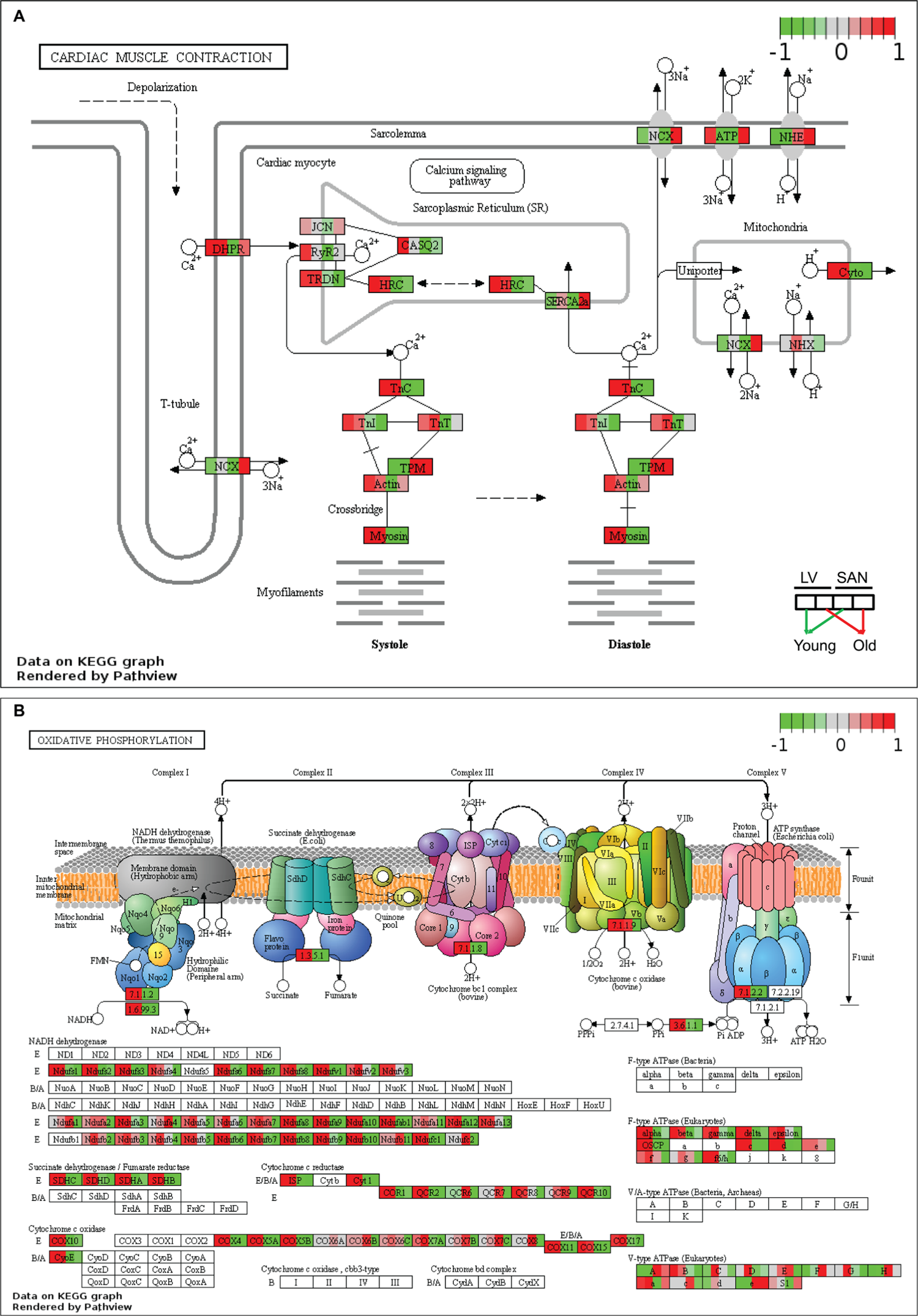
Visualization of KEGG pathways whose genes were expressed more in LVs and in the cluster1 of Figure 1C. The KEGG pathways of cardiac muscle contraction (**A**) and oxidative phosphorylation (**B**) were remodeled by PathView. The color denotes the scaled average FPKM value in each group: from the left to the right, young LV, old LV, young SAN, and old SAN. The redder, the higher scaled value, indicating expressed more in the corresponding group; the greener, the lower scaled value, indicating expressed less in the corresponding group.

#### 3.1.2. Impact of age

Relative to young-LV, genes involved in ion signaling pathways and contraction were downregulated in old-LV, consistent with the age-associated reduction of LV contraction in old mice (Figure 2A). SAN showed upregulation of sodium/hydrogen exchanger (NHE), sodium/calcium exchanger (NCX), and sodium/potassium-transporting ATPase (ATP) genes with aging (Figure 2A). The regulation of those ion exchangers may be linked to the altered electrical impulsive signals in old SAN.

With aging, the expression of some mitochondrial complex and oxidative phosphorylation genes was also changed (Figure 2B). NADH dehydrogenase1 alpha and beta subcomplex subunits, and cytochrome c oxidase subunits, were increased in LV while decreased in SAN. These results point to the age-associated discordant regulation of contraction and mitochondrial function between SAN and LV.

### 3.2. Genes related to post-transcriptional processing

#### 3.2.1. More highly expressed in SAN than LV

As noted in Figure 1C, Cluster2, RNA and protein processing signaling was more positively enriched in SAN vs. LV. After transcription, RNAs experience multiple steps, e.g., splicing, transport, translation, modification, folding, degradation, turnover, etc., in the context of maintenance of the internal homeostasis and normal cellular functions.

Ribosomes execute a major biological function in synthesizing proteins from mRNA in the ribosome. KEGG ribosome pathway showed higher expression of genes in SAN relative to LV (Figure 3A). Protein processing steps beyond translation are also important in the maintenance of protein homeostasis (proteostasis). Intriguingly, the relatively higher level of protein synthesis in SAN vs. LV was accompanied by an increase in the expression of genes involved in protein processing in endoplasmic reticulum (ER) (Figure 1E and 3B). Expression of chaperone (NEF, GRP94, and Hsp40) and reglucosylation (UGGT) genes was also increased in SAN vs. LV. Following ER processing, the correctly folded proteins are transported into Golgi bodies via ERManI, VIP36, Sec family, etc. (grey oval in Figure 3B), and misfolded proteins undergo ER-associated degradation (ERAD), ubiquitination and degradation within the proteasome. The expression of genes within ubiquitin ligase complex and proteasome, however, was not generally higher in SAN than LV (blue irregular box and green circle in Figure 3B). Rather, the KEGG proteasome pathway (Figure 3C) was enriched from the genes in Cluster1 (containing LV-specific genes) (Figures 1E). Misfolded proteins that cannot be corrected or degraded accumulate in the cytoplasm, leading to ER stress. As shown in the Figure 3C bottom left, the unfolded protein response (UPR) genes were more highly expressed in SAN vs. LV, consistent with a triggering of a UPR response in the context of ER stress, and also consistent with an increased rate of protein synthesis in SAN vs. LV.

**Figure 3.**
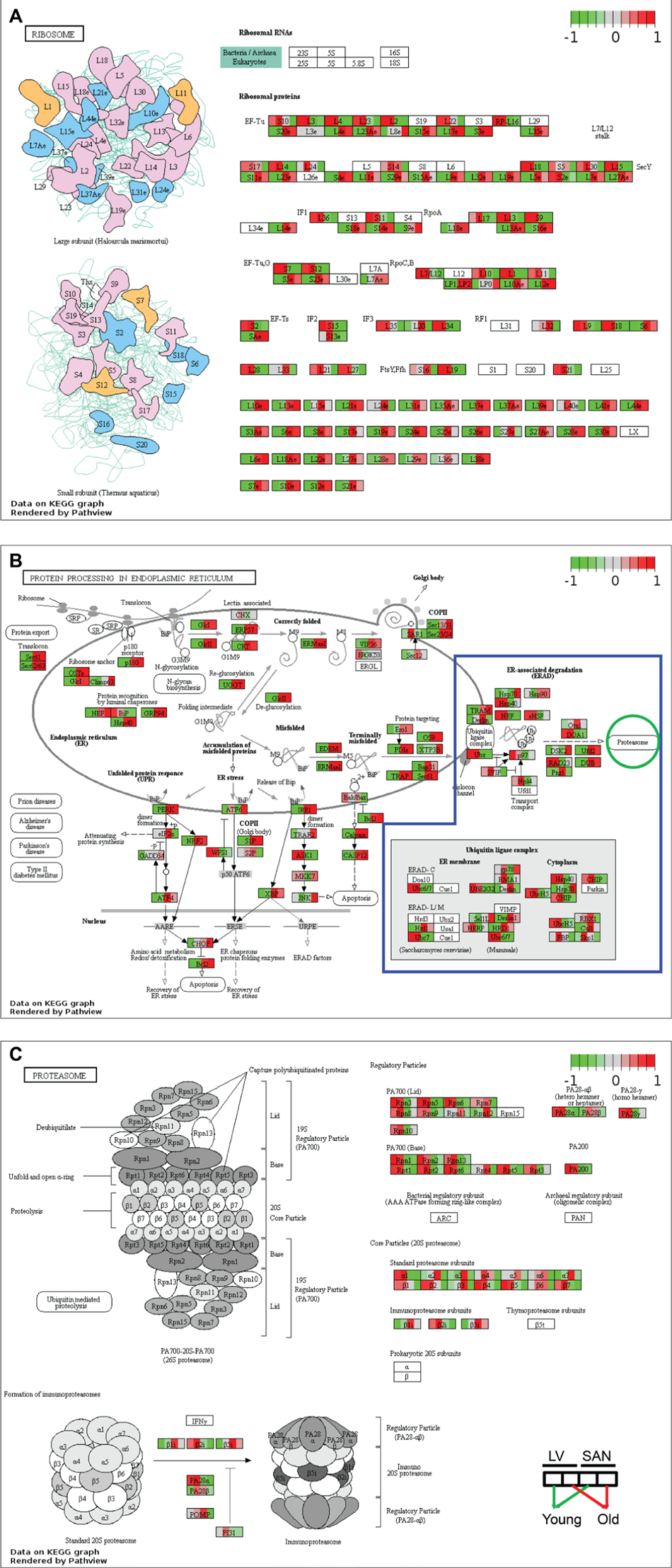
Visualization of KEGG pathways whose genes were expressed more in SANs and in the cluster2 of Figure 1C and proteasome. The KEGG pathways of ribosome (**A**) and protein processing in endoplasmic reticulum (**B**) were remodeled by PathView. In Figure 3B, genes in the blue irregular box are involved in the protein degradation, processing proteins to be degraded in the proteasome (green circle). Then the KEGG pathway of proteasome (**C**) was also remodeled by PathView.

#### 3.2.2. Impact of age

Similar to the age-associated changes in the KEGG pathways of oxidative phosphorylation, many genes within ribosomal complex were upregulated in LV during aging while many ribosomal genes were downregulated with aging in SAN. Because protein synthesis consumes a high level of energy, the concordant upregulation of LV genes points to upregulation in LV protein synthesis with aging; the concordant downregulation of oxidative phosphorylation and ribosomal genes in SAN with aging points to downregulation in SAN protein synthesis with aging. Further, the expression of genes in PA700 (Base) and core particle (20S) was downregulated in old SAN vs. young SAN, suggesting an attenuation of protein degradation in advanced age (Figure 3C). Downregulation of expression of genes involved in post-transcriptional processing (eIF2α, NRF2, ATF4, Bak/Bax, and XBP) in the old SAN vs. young SAN suggest an age-associated disruption of proteostasis.

## 4. Potential sequela of the reprogrammed transcriptome in SAN and LV during aging related to longevity and cardiovascular diseases

### 4.1. Longevity regulating KEGG pathways

The KEGG longevity regulating pathway is conserved across multiple species (Supplementary Figure 4). The KEGG longevity regulating pathway in mammals is depicted in Figure 4 Panel A. In our samples, genes within the KEGG longevity regulating pathway were aggregated in cluster 3 (Figure 1C and 1E). The regulation of longevity is complex, and the KEGG longevity regulating pathway **is linked to numerous other pathways**. The relative expression of **genes** involved in these pathways in SAN and LV was summarized in Figure 4B. In general, the relative expression of genes within insulin, PI3K-Akt signaling, Adenylyl cyclase (AC), SIRT1, FOXO, and autophagy signaling pathways was lower in LV than SAN, whereas the relative expression of genes within AMPK, mTOR, and reactive oxidative species (ROS) scavenging pathways was greater in LV than SAN (Figure 4B). The greater relative expression of genes within insulin signaling and PI3K-Akt pathways within the SAN suggests that insulin signaling and glucose metabolism may be increased in SAN relative to LV (Figure 4B).

**Figure 4.**
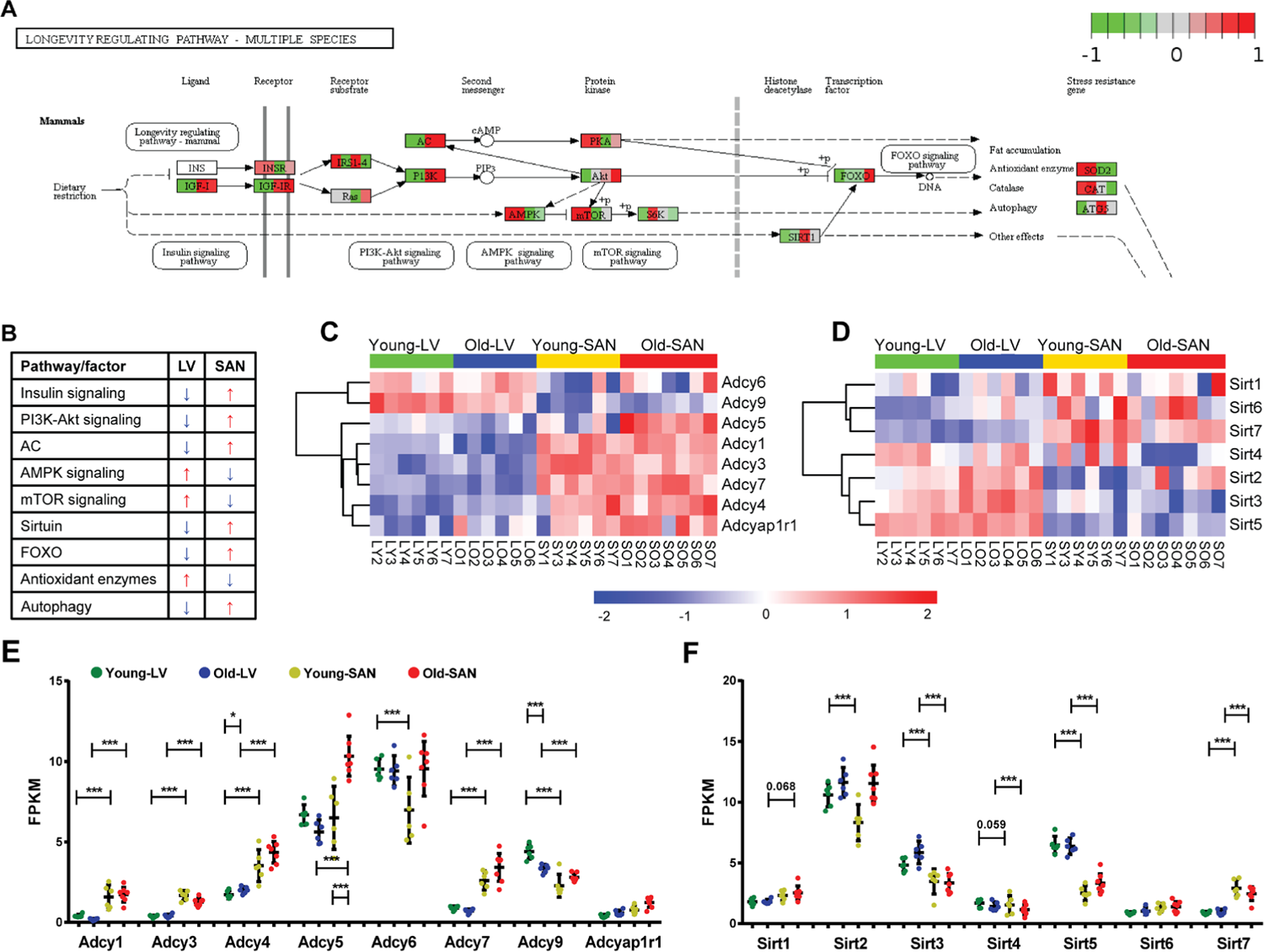
Visualization of the mammals-specific longevity regulating pathway whose genes were clustered in the cluster3 of Figure 1C. **A.** The mammals-specific longevity regulating pathway was remodeled by PathView. **B.** The relative gene expressions were summarized. The blue arrow denotes the lower expression, and the red arrow denotes the higher expression. **C-D.** Heatmap of adenylyl cyclase genes (**C**) and sirtuin genes (**D**) in the four groups. **E-F.** Scatter plot of adenylyl cyclase genes (**E**) and sirtuin genes (**F**) with the q-values between two groups: *, q-value < 0.05; **, q-value < 0.01; ***, q-value < 0.001. The bar shows the mean and 95% CI.

#### 4.1.1. mTOR signaling

Expression of genes within mTOR signaling pathway was slightly increased in LV than SAN. A higher level of mTOR signaling in LV vs. SAN (Figure 4B) may be linked to the less efficient autophagy in LV vs. SAN.^14^ Of note, however, in advanced age, ATG5, an important inducer of autophagy, is reduced in SAN (Figure 4A).

#### 4.1.2. AMPK signaling

Same to mTOR, expression of genes within AMPK signaling pathway that counteracts some facets of mTOR signaling^15^ was also higher in LV vs. SAN (Figure 4A and 4B). These chamber differences in mTOR and AMPK signaling is likely linked to differences in LV vs. SAN with respect to enrichments in ribosome and protein processing, mitochondria, and anti-oxidant defenses in LV vs. SAN (Figure 4A). There were no evident changes in AMPK signaling in LV or SAN with age.

#### 4.1.3. Sirtuin family

Expression of Sirt1, Sirt6, Sirt7 and Foxo, potential prolongevity factors,^16, 17^ was relatively higher in SAN than LV. Expression of Sirt3 and Sirt5, which promote defence against oxidative stress,^18, 19^ was increased in LV vs. SAN, consistent with gene expression patterns of oxidative phosphorylation and antioxidant enzymes (Figure 4D and 4F).

Of note, although Foxo was increased with age in SAN, Sirt1, Sirt6 or Sirt7 were not increased with age in SAN. Interestingly, Sirt2, which protects against pathological cardiac hypertrophy,^20^ was increased with age in LV and SAN, whereas Sirt4, which is linked to pathologic cardiac hypertrophy,^21^ was downregulated with age in LV and SAN (Figure 4D and 4F), suggesting that LV and SAN upregulated Sirt2 but downregulated Sirt4 to fight against cardiac stress during aging. Expression of Sirt3 was also increased with age in LV.

#### 4.1.4. Adenylyl cyclase (AC) signaling

Adenylyl cyclase (AC) signaling, which is very important in regulation of cardiac function,^22, 23^ is also a component of longevity regulation signaling pathway (Figure 4A). A heatmap of the expressions of genes within the family of AC, which catalyze the conversion of adenosine triphosphate (ATP) to 3’,5’-cyclic AMP (cAMP) and pyrophosphate,^24^ is shown in Figure 4C, and the FPKM values of ACs in young and old, LV and SAN are compared in the scatter plot in Figure 4E. Except for Adcy6 and Adcy9, other members in the AC family were expressed more in SAN than in LV, consistent with the important role of AC-cAMP-PKA axis in initiating and executing the pacemaker signals in SAN.^25, 26^ There were no evident changes in AC signaling in LV or SAN with age.

### 4.2. Prolongevity and antilongevity genes recorded and curated in the GenAge database

In addition to analyzing our RNA-seq data within longevity related KEGG pathways, we also analyzed the expression pattern of aging-related genes in GenAge, an aging gene database that distinguishes prolongevity from antilongevity genes.^27^

#### 4.2.1. SAN vs. LV

As shown in Figure 5A and 5B, the majority of prolongevity genes were more highly expressed in SAN than LV, while a minority of antilongevity genes more highly expressed in SAN than LV. This is consistent with the idea that the organism prioritizes gene expression to preserve the life of its SAN.

**Figure 5.**
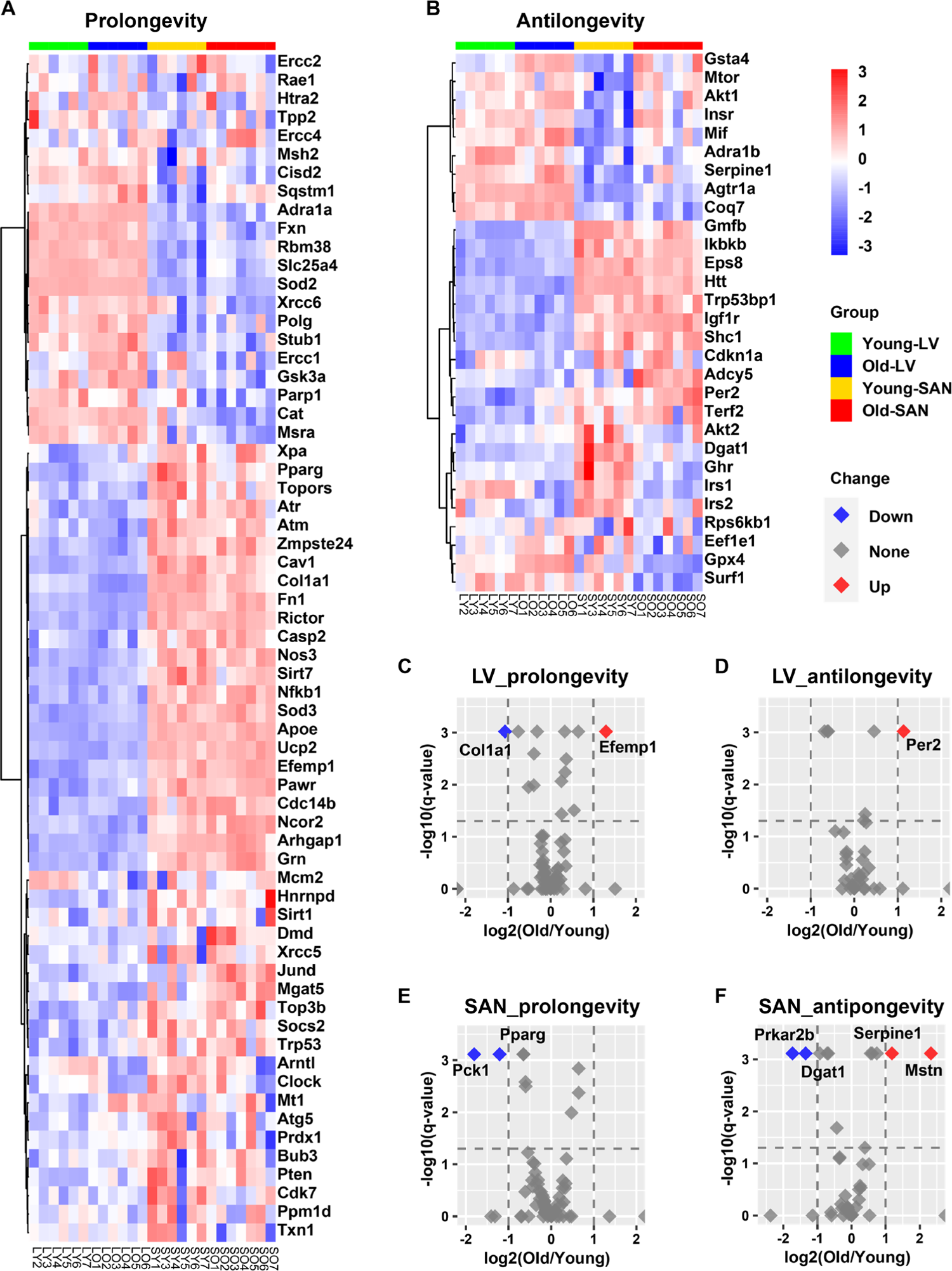
Longevity effect analysis of RNA-seq data in this project. **A-B.** Heatmap of possible prolongevity (**A**) and antilongevity (**B**) genes by mapping our RNA-seq data to the GenAge database. **C-F.** Volcano plots of possible prolongevity (**C and E**) and antilongevity (**D and F**) genes in LV (**C** and **D**) and SAN (**E** and **F**). The cutoffs of the regulation are: (1) -log10(q-value) = 1.3, i.e., q-value = 0.05, and (2) absolute value of log2(Old/Young) = 1. The red dot (Up) denotes the upregulated gene in old tissues; blue (Down) denotes downregulation; grey (None) denotes non-significant.

#### 4.2.2. Impact of age

In SAN, two prolongevity genes, phosphoenolpyruvate carboxykinase 1 (Pck1)^28^ and peroxisome proliferator activated receptor gamma (Pparg),^29, 30^ were downregulated with aging; and four antilongevity genes were changed with aging: protein kinase, cAMP dependent regulatory, type II beta (Prkar2b)^31^ and diacylglycerol O-acyltransferase 1 (Dgat1)^32^ were downregulated, while serine (or cysteine) peptidase inhibitor, clade E, member 1 (Serpine1; also known as plasminogen activator inhibitor-1 (PAI-1))^33^ and myostatin (Mstn; also known as growth differentiation factor 8 (GDF-8))^34^ were upregulated (Figure 5E and 5F).

In LV, two prolongevity genes were changed with aging: collagen, type I, alpha 1 (Col1a1)^35^ was downregulated and epidermal growth factor-containing fibulin-like extracellular matrix protein 1 (Efemp1; all known as fibulin-3)^36^ was upregulated. It’s interesting to note that fibroses increases with aging while elastin becomes degraded with aging. One LV antilongevity gene, period circadian clock 2 (Per2)^37^ was upregulated (Figure 5C and 5D).

### 4.3. Cardiovascular disease marker genes

Because advanced age is the major cause of cardiovascular diseases (CVDs),^2–4^ we mapped a group of cardiovascular disease marker genes^38^ to the transcriptome of our samples. In general, the marker gene expression of abdominal aortic aneurysm, coronary artery disease, heart failure, hypertension, insulin resistance, myocardial infarction, stroke, and vascular disease was relatively higher in SAN, whereas that of cardiomyopathy, dyslipidemia, left ventricular noncompaction, and septal hypertrophy was higher in LV v. SAN (Supplementary Figure 5). This suggested that differential changes in gene expression within the LV and SAN contribute to various CVDs.

Importantly, expression of genes related to CVDs tended to include most of the genes showing a trend to increase during aging in both LV and SAN (Supplementary Figure 5). This suggests the impaired LV and SAN functions are associated with increased incidence of CVDs in advanced age.

## 5. Upstream transcriptional and post-transcriptional regulators

### 5.1. Transcriptional regulators

#### 5.1.1. Chamber-specific transcriptional regulators

Because gene expression is usually under the control of many transcriptional regulators, we employed two tools and methods to predict the upstream transcription regulators. First, we performed the epigenetic **L**andscape **I**n **S**ilico deletion **A**nalysis (LISA)^12^ to predict the transcription regulators that were likely to be regulators of gene expression in LV (bottom right in Figure 6A and 6B) or in SAN (top left in Figure 6A and 6B). Within the ChIP-seq model, TBX and NKX2-5 were predicted to regulate LV gene expression, while CCAAT/enhancer-binding protein alpha (CEBPA), transcription factor PU.1 (SPI1), and Wilms’ tumor protein (WT1) were predicted to regulate SAN gene expression (Figure 6A).

**Figure 6.**
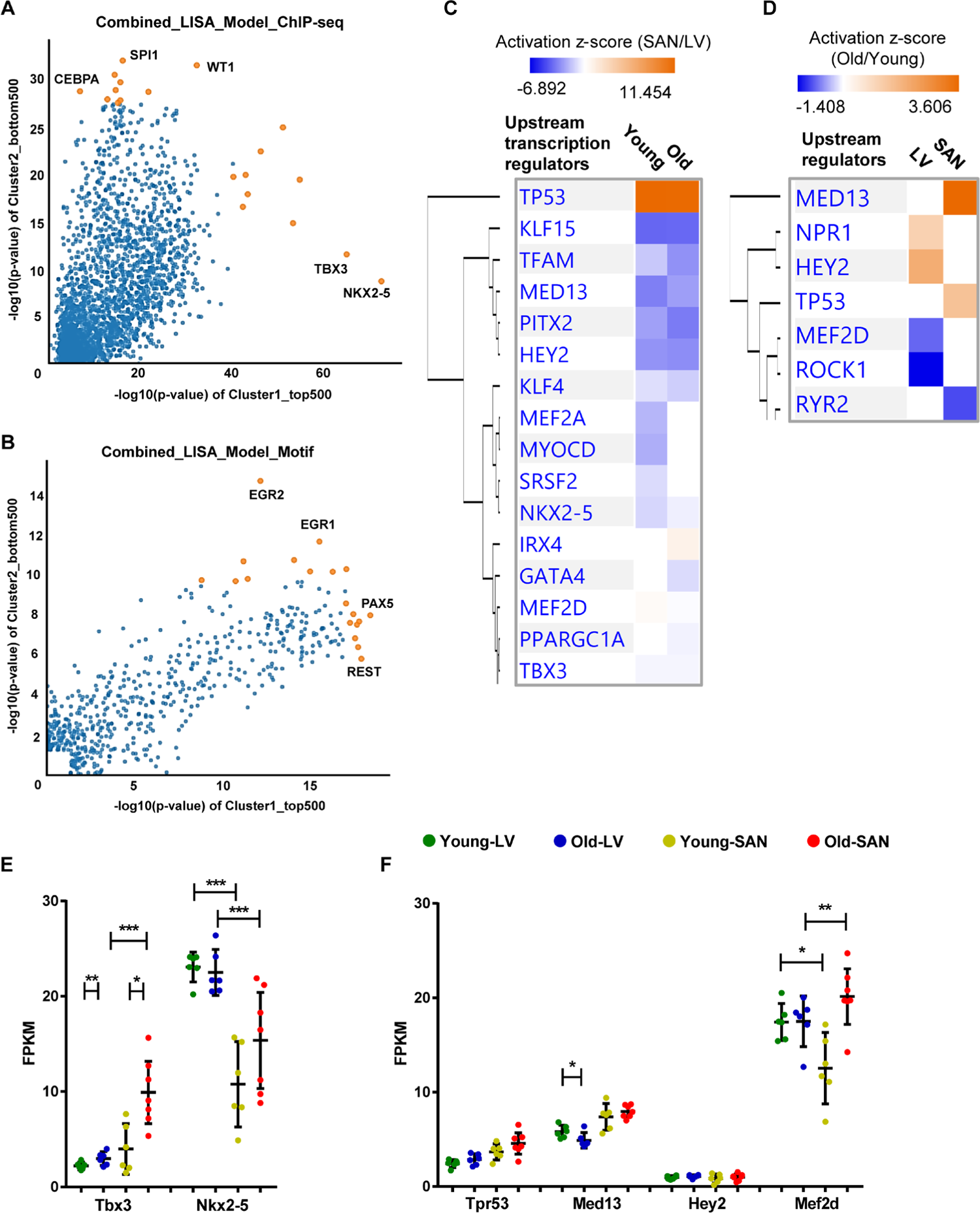
Upstream transcription regulator analysis. **A-B.** Prediction of upstream transcription regulators by the comparison of top 500 genes in Cluster1 and bottom 500 genes in Cluster2 in Figure 1C, using the ChIP-seq (**A**) and Motif (**B**) model of LISA database. **C-D.** Comparison of the upstream regulators predicted in the IPA software by the comparison of SAN/LV (**C**) and Old/Young (**D**). **E-F.** Scatter plots of transcription regulators genes. Transcription regulators consistently predicted using LISA (**A** and **B**) and IPA (**C**) are displayed in Figure **E**. Transcription regulators predicted by the comparison of SAN/LV (**C**) and Old/Young (**D**) are displayed in Figure **F**. The bar shows the mean and 95% CI. q-values between two groups: *, q-value < 0.05; **, q-value < 0.01; ***, q-value < 0.001.

In the motif model, paired box protein Pax-5 (PAX5) and RE1-Silencing Transcription factor (REST, also known as Neuron-Restrictive Silencer Factor (NRSF)) were predicted to regulate LV gene expression, while two early growth response proteins (EGR1 and EGR2) were predicted to regulate SAN gene expression (Figure 6B).

Next, in IPA, NKX2-5 and TBX3 were also predicted as the upstream transcription regulators that were more activated in LV relative to SAN (blue color in Figure 6C). A comparison of regulator activity and gene expression indicated that changes in the gene expression pattern of NKX2-5 but not that of TBX3 was in line with their predicted activity (Figure 6E), indicating that the higher expression of NKX2-5 induced the formation of LV-specific gene expression pattern.

Four transcription regulators, TP53, MED13, HEY2 and MEF2D, that were predicted (Figure 6C and 6D) were also identified in the transcriptome (Figure 6F). TP53 (gene: Tpr53) was more activated in SAN vs. LV (Figure 6C). Mediator complex subunit 13 (MED13), Hairy/enhancer-of-split related with YRPW motif protein 2 (HEY2, also known as cardiovascular helix-loop-helix factor 1 (CHF1)), and Myocyte-specific enhancer factor 2D (MEF2D), were more activated in LV relative to SAN (Figure 6C).

#### 5.1.2. Age-associated transcriptional regulators

In addition to the above prediction based on the differentially expressed genes between chambers, we also used the IPA to predict the age-associated changes in transcription regulators (Figure 6D). Of the four transcription regulators, TP53 (gene: Tpr53) activity was enhanced only in old vs. young SAN (Figure 6C and 6D). The gene expression of MED13, which promotes systemic energy expenditure within the heart,^39^ was reduced in old LV (Figure 6F) but its transcriptional activity was activated in SAN during aging (Figure 6D), suggesting the disrupted metabolism in the heart during aging. The activity of HEY2, a negative transcription regulator of cardiac myocytes,^40^ was increased in old LV, suggesting it may contribute to the repressed number and function of cardiomyocytes (Figure 6D). The activity of MEF2D, which plays a role in stress-dependent cardiac remodeling,^41^ was reduced in Old vs. Young LV and its gene expression was higher in old mice (SAN/LV) may underlie the remodeled heart in advanced age (Figure 6D and 6F).

### 5.2. Post-transcriptional regulators

#### 5.2.1. pre-miRNAs

In addition to transcriptional regulation, post-transcriptional regulation is also an important determinant of mRNA and protein levels. In this transcriptome, we identified many pre-miRNAs and mRNAs in our analysis. Expression of pre-miRNAs having z-scores greater than zero in at least three samples in each group were displayed in heatmap (Figure 7A). We first compared pre-miRNAs between chambers (SAN/LV) (left or right pie in Figure 7B separately). In accord with the heatmap in Figure 7A, expression of nearly all significantly differentially expressed pre-miRNAs (q-value <= 0.1) was greater (log2_FC > 0) in SAN than LV regardless of age. Particularly, expression of two pre-miRNAs, Mir5136 and Mir770, was higher in SAN than LV in both young and old mice (Figure 7B).

**Figure 7.**
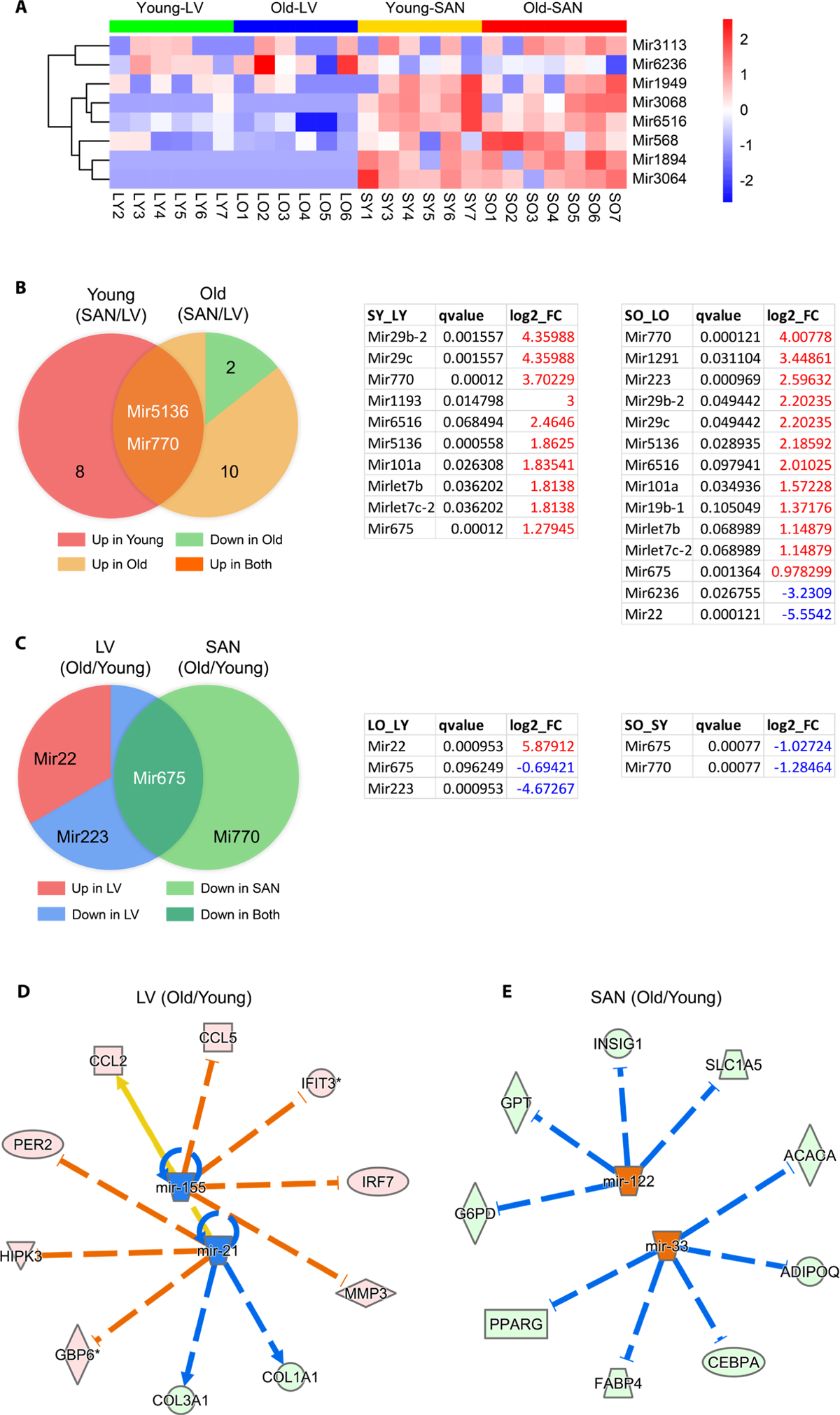
microRNA analysis. **A.** Heatmap of pre-miRNAs which were expressed in >= 3 samples in each group. **B-C.** Venn diagrams (left) and corresponding tables (right) of differentially expressed genes. **D-E.** Regulation of miRNAs on differentially expressed genes between old and young samples. Genes identified in the RNA-seq are distributed in the circle, and predicted upstream mature miRNAs are in the center. The pink dot denotes the upregulated gene in old tissues, the light green dot denotes the downregulated gene, the orange dot denotes the predicted increased miRNA, and the blue dot demotes the predicted decreased miRNA. The arrow denotes activation, and the line dotted line denotes inhibition. The orange and blue edges suggest that the gene expression changes were in accord with the predicted function of the miRNAs. If the downstream gene is upregulated, the edge is in orange; downregulation, in blue. The yellow edge suggests the discordant function.

By consideration of age, two pre-miRNAs, Mir6236 and Mir22, were expressed fewer in SAN vs. LV only in old mice (Figure 7B). Regardless of chamber type, there were more pre-miRNAs whose expression was downregulated in old vs. young samples (Figure 7C). Both LV and SAN Mir675 expression was downregulated with aging (Figure 7C), but only Mir22 expression was higher in old vs. young LV (Figure 7C).

#### 5.2.2. Mature miRNAs

Because the processed mature miRNA functions as the post-transcriptional regulator of mRNA stability and translation,^42^ we continued to analyze the mature miRNA in IPA. Based on the four pairs of comparison (LO vs. LY, SO vs. SY, SY vs. LT, and SO vs. LO), two, two, five and four upstream mature miRNAs, respectively, were predicted to be significantly inhibited or activated (absolute(z-score) >= 2).

Regardless of age, the predicted miRNAs displayed lower activities in SAN vs. LV, in both young (Supplementary Figure 6) and old (Supplementary Figure 6) mice. With respect to aging, mir-21 and mir-155 were predicted to be inhibited in old vs. young LV (Figure 7D), and the inhibition of mir-21 accounted for the decreased expression of Col1a1 (Figure 5C) and increased expression of Per2 (Figure 5D). Mir-122 and mir-33, which regulate many metabolic genes, were predicted to be activated in old vs. young SAN to (Figure 7E). For example, the activation of mir-33 accounted for the decreased expression of Pparg in old SAN (Figure 5E).

## Discussion

The heart is composed of different specialized functional parts.^5–7^ The SAN (major location of intrinsic pace maker cells) and LV (consisting of large number of working cardiomyocytes) are quite distinct in term of structure and function,^6, 9^ but their different molecular features have never been compared in omics scale, not to mention in the context of aging, which is a detrimental factor in the development of CVDs.^1^ To our knowledge, this project is the first transcriptomic analysis in which LV and SAN were compared with respect to not only age but also chamber-specific differences as well as differences in direction of age-associated changes by heart chamber. Specifically, we compared expression patterns across the four groups of heart tissues (LY, LO, SY and SO). We combined multiple bioinformatics analyses and selected experimental validations to reveal predicted upstream regulators and downstream effects within cardiac transcriptome of the four groups in order to deduce possible molecular mechanisms in SAN vs. LV, while taking into account of the effect of age.

We observed that changes in gene expression and signaling pathways that are implicated in cardiac functions, not only differed by chamber, but also differed by age, often in a chamber-specific manner. Differentially altered pathways, including mitochondrial function, metabolism, RNA and protein processing, stress response signaling, etc., and upstream regulators, including transcription regulators and miRNAs, which could be regarded as potential therapeutic targets to enhance cardiac function.

## 1. Difference between chambers (SAN and LV)

According to hierarchical analysis and PCA, differences between chambers (SAN vs. LV) were greater than differences by age (Figure 1A and 1B).

### 1.1. Gene level

Heatmap clustering identified three clusters of genes with a similar number of genes in Cluster1 (containing LV-specific genes) or in Cluster2 (containing SAN-specific genes) (Figure 1C). These clusters included, but were not limited to, some well-acknowledged, cardiac-specific genes (Supplementary Figure 2).

### 1.2. Enrichment level

LV or SAN-specific genes were differentially enriched in GO terms and KEGG pathways (Figure 1 and Supplementary Figure 3). Given that four groups were simultaneously compared in our analysis, it is important to note that the functional differences mainly at the chamber level could occur without respect to age.

Specifically, oxidative phosphorylation and metabolism related terms and pathways were enriched in Cluster1 (containing LV-specific genes), while RNA processing, protein processing, and signaling regulation related terms and pathways were enriched in Cluster2 (containing SAN-specific genes) (Figure 1D and 1E). At the pathway activity level, many metabolic pathways, especially those related to energy metabolism in mitochondria, consistently displayed higher activity in LV vs. SAN (Figure 1F). Mitochondrial metabolism produces a large amount of energy required for LV contraction and ejection of blood from heart. For instance, the activity of fatty acid β-oxidation pathway, the major source of energy production in heart, was greater in LV vs. SAN. Commensurate with activated oxidative phosphorylation, oxidative stress response signaling (Figure 1F) and antioxidant enzyme genes (Figure 4B) were also induced to a greater extend in LV than SAN. This is not surprising because enhanced oxidative phosphorylation would be expected to generate enhanced reactive oxidative species (ROS).

Many other crucial signaling pathways in cardiac function also differed between SAN and LV. Because beta adrenergic receptor signaling activates adenylyl cyclase (AC) signaling to increase heart rate via effects within the SAN and also increase LV contraction, we focused on the tissue difference in AC signaling (Figure 4B). Except for Adcy6 and Adcy9, other members in the AC family were expressed more in SAN than in LV, consistent with the important role of AC-cAMP-PKA axis in the coupled-clock system that initiates impulse within the SAN that determines the beating rhythm of the heart.^25, 26^

mTOR signaling is linked to many cell processes that are necessary for maintenance of normal heart structure and function, protein synthesis, proteins processing, mitochondria, antioxidant, etc.^43–46^ Expressions of genes within mTOR signaling pathway were slightly more increased in LV than SAN. Similar to mTOR, expression of genes within AMPK signaling pathway was also higher in LV vs. SAN. This difference in AMPK signaling between chambers is concordant with the difference of mTOR signaling within chambers (Figure 3A and 3B, Figure 4A). AMPK signaling counteracts some facets of mTOR signaling via: (1) inhibition of mTORC1 and protein synthesis,^45^ concordant with the lower gene expressions of ribosome and protein processing in LV (Figure 3A and 3B); (2) promotion of gene transcription in mitochondria,^46^ contributing to the enrichments of mitochondria in LV (Figure 1 and Supplementary Figure 3); and (3) activation of anti-oxidant defenses,^46^ in line with the higher level of antioxidant enzymes (SOD2 and CAT) (Figure 4A and 4B) required to clear the excess ROS as a side product of mitochondrial respiration (Figure 1 and Supplementary Figure 3). Sirtuins are class III deacetylases, implicated in influencing metabolism and stress resistance.^47, 48^ The four members existing in mitochondria (Sirt3, Sirt4, and Sirt5) and another stress responder outside of mitochondria (Sirt2) were expressed more in LV (Figure 4D).

### 1.3. Regulator level

The transcription factor, NKX2-5, was predicted to be more highly activated in LV than SAN, whereas TP53 was predicted to be activated more in SAN than LV (Figure 6). miRNAs are post-transcriptional regulators of gene expression. Intriguingly, expression of those miRNAs pre-mature forms and mature form activities displayed the contrasting trends in SAN vs. LV (Figure 7, and Supplementary Figure 6 and 7), which may result from chamber-specific differences in miRNA processing during maturation. Such differential regulation of gene expression may result in the different contributions of specific genes to age-associated changes and pathophysiology related to CVDs in SAN vs. LV (Figure 5 and Supplementary Figure 5).

## 2. Impact of age

Our results revealed that age-associated differences occurred in expression and regulation of many cardiac genes, without reference to chamber. For example, Mir675 was downregulated in *both* LV and SAN in old vs. young samples (Figure 7C). Although some marker genes associated with CVDs were in more highly expressed in either LV or SAN, most CVD maker genes were upregulated in *both* LV and SAN in old vs. young samples (Supplementary Figure 5), suggesting that these genes may be associated with increased morbidity and mortality as age advances. An association of CVD marker genes with increased morbidity and mortality may be related to either the increase of antilongevity genes observed in old vs. young LV (Figure 5D) or the reduction of prolongevity genes observed in old vs. young SAN (Figure 5E).

## 3. Chamber-specific impact of age

Despite some similar age-associated changes in gene expression in both LV and SAN, age was also associated with chamber-specific differences. Specifically, genes involved in GO terms, KEGG pathways and canonical pathways signaling were under different control in old vs. young in LV or SAN (Figure 1 and Supplementary Figure 3). For example, KEGG pathways of cardiac muscle contraction and oxidative phosphorylation were enriched in genes in Cluster1 (containing LV-specific genes), some genes with these pathways manifested different age-associated changes in LV or SAN. Examples included genes involved in calcium signaling, ion exchangers, NADH dehydrogenase1, cytochrome c oxidase, ribosome, protein processing, etc. (Figure 2 and 3). Discordant regulation of contraction and mitochondrial function genes between SAN and LV with aging might be implicated in chamber-specific disease manifestations with aging, e.g., chronic LV contraction failure vs. sick sinus syndrome.^1^

Upstream regulators of gene expression had differential patterns in SAN vs. LV in old vs. young samples. For instance, in LV, NPR1 and HEY2 were activated while MEF2D and ROCK1 were inhibited; in SAN, MED13 and TP53 were activated while RYR2 was inhibited (Figure 6D and 6F). Pre-miRNAs were significantly differentially regulated in chamber-specific fashion in old vs. young samples (Figure 7C). Additionally, activity of mature miRNAs was differentially impacted by aging in LV or SAN respectively (Figure 7D and 7E). This differential regulation is likely implicated in differential changes in differential changes observed in the longevity regulating pathway (Figure 4) and prolongevity and antilongevity gene expression profiles (Figure 5). Therefore, such discordant alterations in LV and SAN with aging may disrupt the functional coherence between chambers observed in CVD progression as age advances.

## Limitation

Our analysis based on deep RNA-seq and bioinformatics analysis uncovered many novel aspects of chamber-specific and age-associated differences. Specifically, we identified a large number of differentially expressed genes, including mRNAs and pre-miRNAs, among the four groups (LY, LO, SY and SO), enriched and predicted important downstream signaling pathways, and predicted upstream regulators. Our findings regarding longevity regulating pathway and longevity gene expression might be leveraged to improve cardiac health even in advanced age. However, this discovery regarding gene expression and activities of pathways and regulators remained to be verified. Future studies are required to distinguish the actual gene expression patterns and predicted activity in the four groups and the discordance between pre-miRNA expression and mature miRNA activity results from multiple miRNA processing steps, e.g., splicing, trafficking, degradation, etc.

## Acknowledgements

We thank all members in Laboratory of Cardiovascular Science, National Institute on Aging, National Institutes of Health, for technical assistance and discussion. This work utilized the computational resources of the NIH HPC Biowulf cluster (http://hpc.nih.gov).

## Sources of Funding

Intramural Research Program at National Institute on Aging, National Institutes of Health.

## Disclosures

None.

---

**Supplementary Figure 1.**
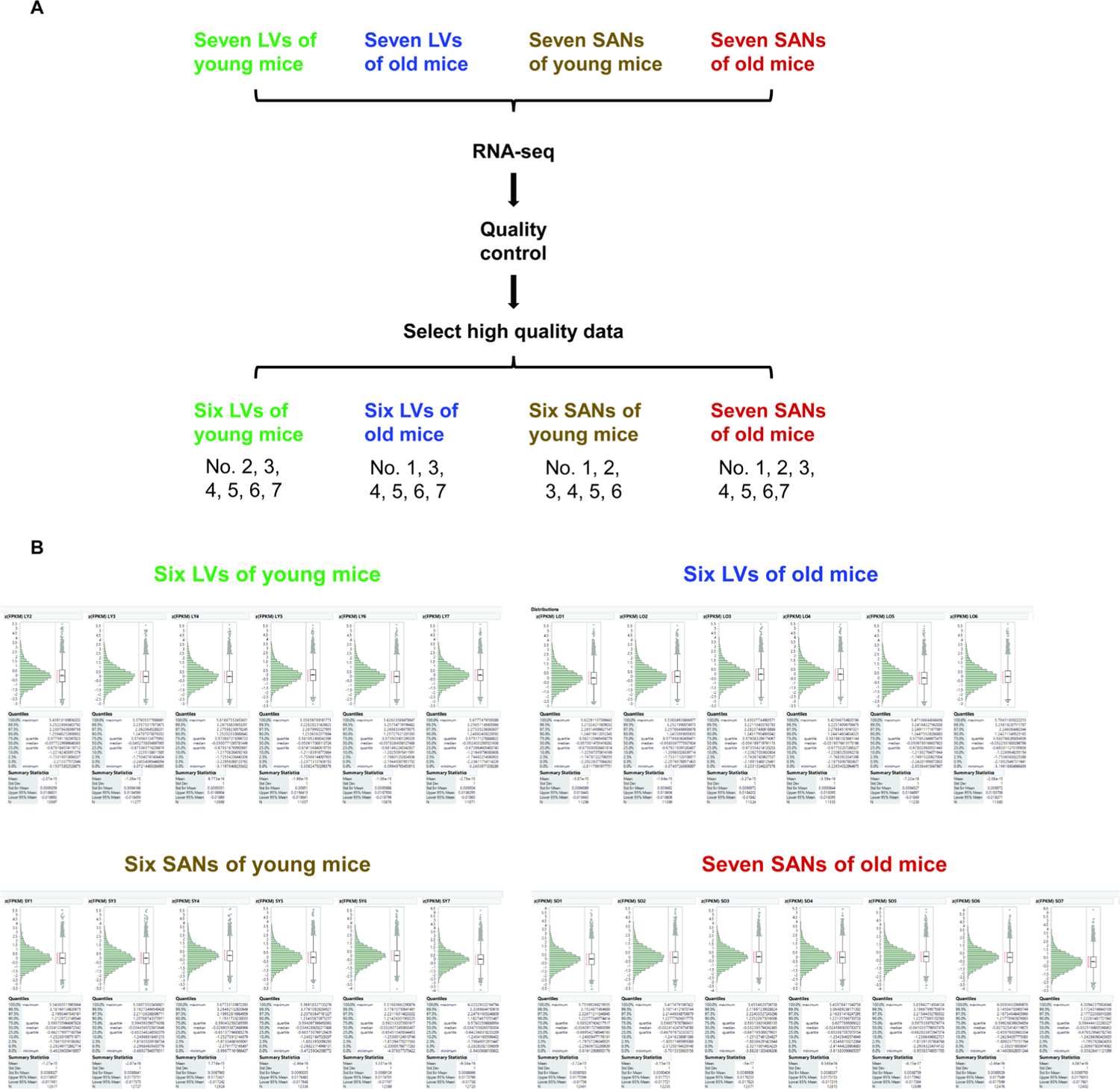
Four groups of samples and the distribution of their FPKM values. **A.** Four groups of samples were included in this project: left ventricles (LVs) of young mice (LY), LVs of old mice (LO), sinoatrial nodes (SANs) of young mice (SY), and SANs of old mice (SO). After quality control, six samples in the LY group, six samples in the LO group, six samples in the SY group, and seven samples in the SO group were remained for subsequent analyses. **B.** The FPKM of each sample was z-score normalized to be in the approximate normal distribution.

**Supplementary Figure 2.**
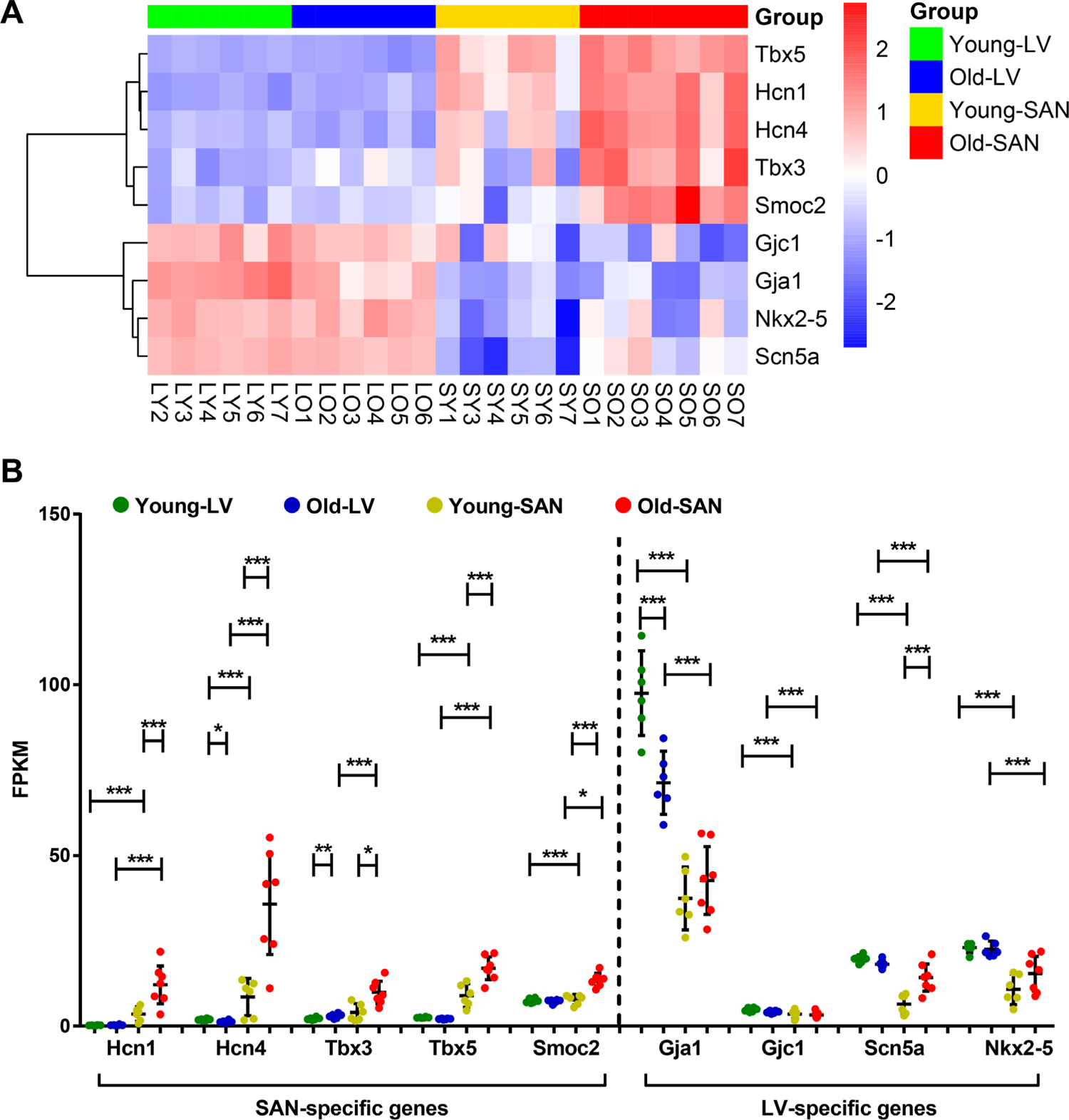
Heart tissue marker genes. **A.** Heatmap of several heart tissue marker genes. **B.** Scatter plot of those heart tissue marker genes with the q-values between two groups: *, q-value < 0.05; **, q-value < 0.01; ***, q-value < 0.001. The bar shows the mean and 95% CI.

**Supplementary Figure 3.**
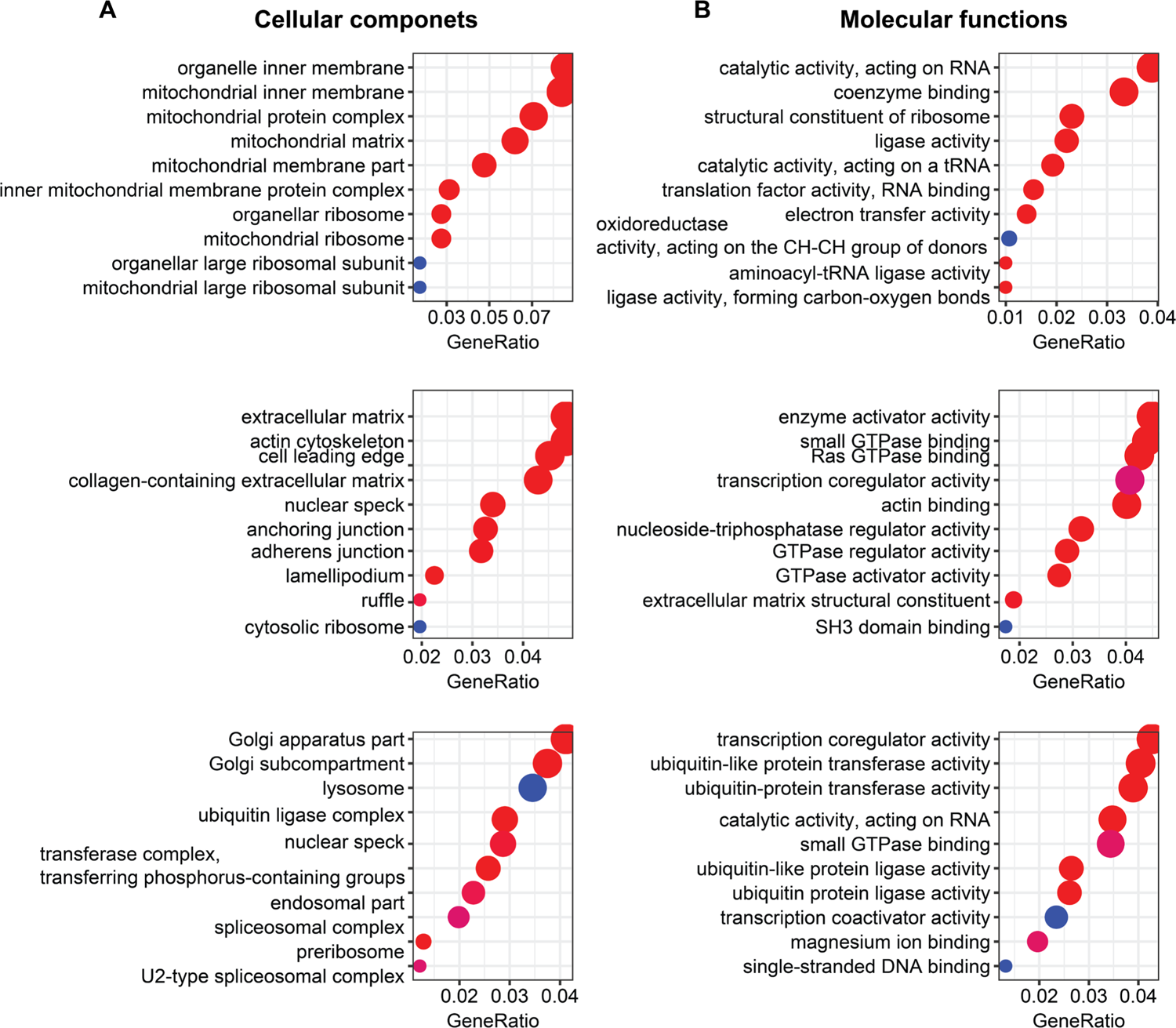
GO analyses with genes in each cluster of Figure 1C. Bubble charts of GO cellular component (**A**) and molecular function (**B**) analysis with genes in each cluster. Same to Figure 1D and 1E, the larger the bubble, the more identified genes in the term. The redder, the less adjusted p-value, indicating more significant.

**Supplementary Figure 4.**
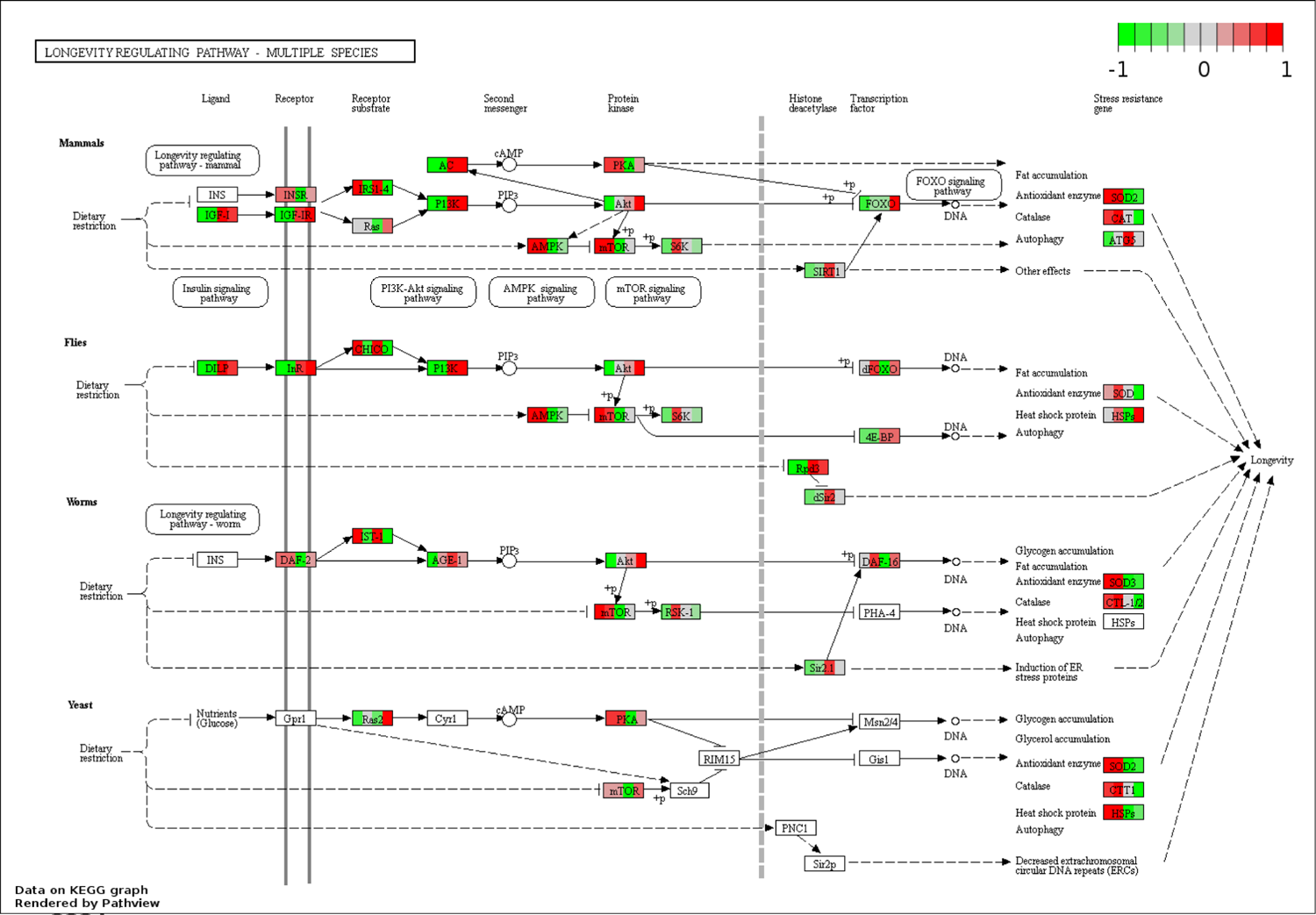
Visualization of the entire longevity regulating pathway (multiple species) whose genes were clustered in the downmost cluster of Figure 1C. The longevity regulating pathway was visualized and compared in multiple species, including mammals, flies, worms and yeast.

**Supplementary Figure 5.**
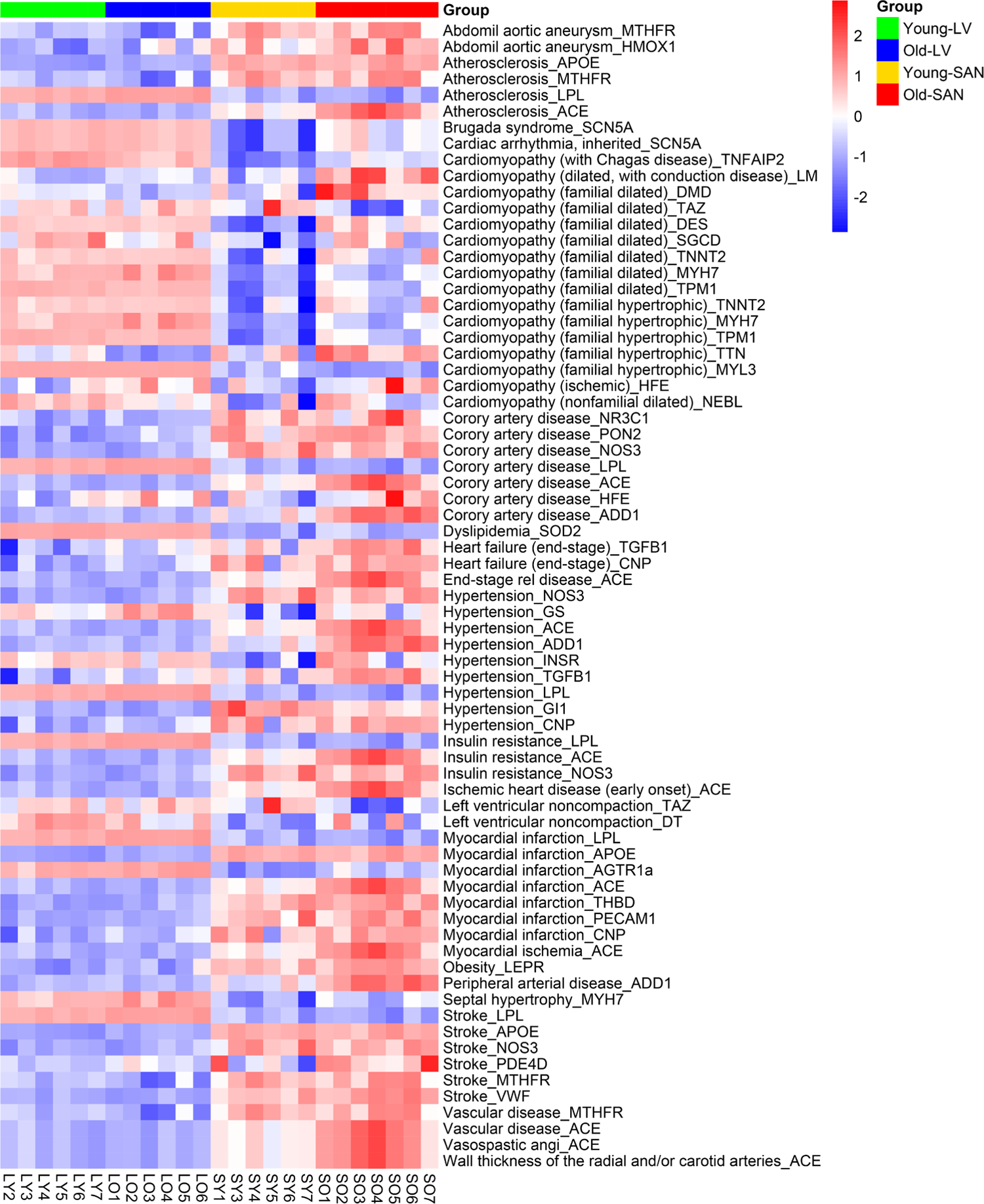
Cardiovascular disease marker genes analysis. Heatmap of cardiovascular disease marker genes.

**Supplementary Figure 6.**
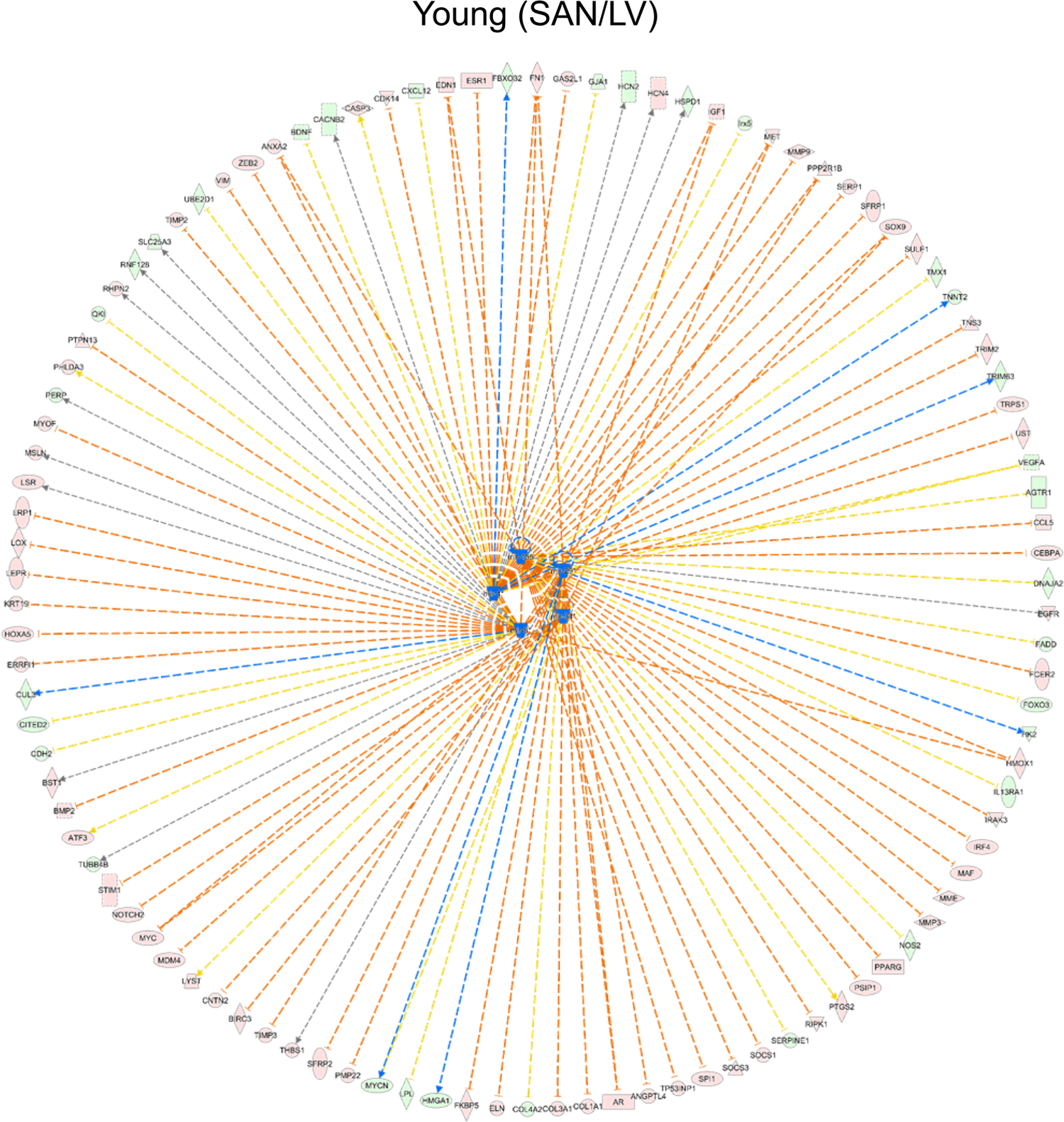
Regulation of miRNAs on differentially expressed genes between SAN and LV tissues in the young mice. Genes identified in the RNA-seq are distributed in the circle, and predicted upstream mature miRNAs, mir-1, mir-155, mir-29, mir-34 and mir-155, are in the center.

**Supplementary Figure 7.**
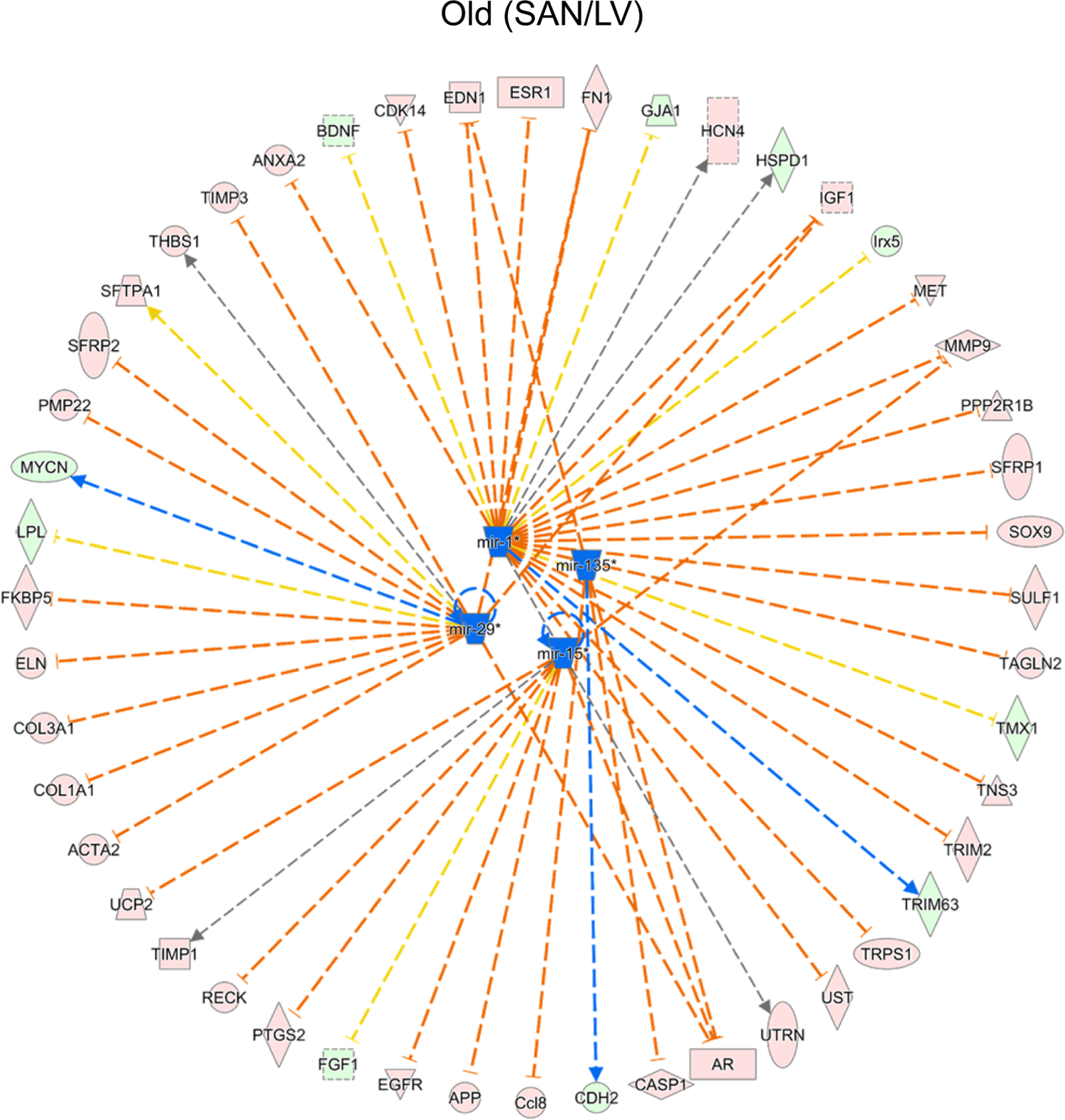
Regulation of miRNAs on differentially expressed genes between SAN and LV tissues in the old mice. Genes identified in the RNA-seq are distributed in the circle, and predicted upstream mature miRNAs, mir-1, mir-135 and mir-15, mir-29, are in the center.

## Notes

### Competing Interest Statement

The authors have declared no competing interest.

